# The ENCODE Imputation Challenge: A critical assessment of methods for cross-cell type imputation of epigenomic profiles

**DOI:** 10.1101/2022.07.30.502157

**Authors:** Jacob Schreiber, Carles Boix, Jin wook Lee, Hongyang Li, Yuanfang Guan, Chun-Chieh Chang, Jen-Chien Chang, Alex Hawkins-Hooker, Bernhard Schölkopf, Gabriele Schweikert, Mateo Rojas Carulla, Arif Canakoglu, Francesco Guzzo, Luca Nanni, Marco Masseroli, Mark James Carman, Pietro Pinoli, Chenyang Hong, Kevin Y. Yip, Jeffrey P. Spence, Sanjit Singh Batra, Yun S. Song, Shaun Mahony, Zheng Zhang, Wuwei Tan, Yang Shen, Yuanfei Sun, Minyi Shi, Jessika Adrian, Richard Sandstrom, Nina Farrell, Jessica Halow, Kristen Lee, Lixia Jiang, Xinqiong Yang, Charles Epstein, J. Seth Strattan, Michael Snyder, Manolis Kellis, William Stafford Noble, Anshul Kundaje, ENCODE Imputation Challenge Participants

## Abstract

Functional genomics experiments are invaluable for understanding mechanisms of gene regulation. However, comprehensively performing all such experiments, even across a fixed set of sample and assay types, is often infeasible in practice. A promising alternative to performing experiments exhaustively is to, instead, perform a core set of experiments and subsequently use machine learning methods to impute the remaining experiments. However, questions remain as to the quality of the imputations, the best approaches for performing imputations, and even what performance measures meaningfully evaluate performance of such models. In this work, we address these questions by comprehensively analyzing imputations from 23 imputation models submitted to the ENCODE Imputation Challenge. We find that measuring the quality of imputations is significantly more challenging than reported in the literature, and is confounded by three factors: major distributional shifts that arise because of differences in data collection and processing over time, the amount of available data per cell type, and redundancy among performance measures. Our systematic analyses suggest several steps that are necessary, but also simple, for fairly evaluating the performance of such models, as well as promising directions for more robust research in this area.

## 1 Introduction

Since their development, high-throughput chromatin profiling assays such as histone ChIP-seq, DNase-seq and ATAC-seq have proven crucial for deciphering gene regulatory elements and characterizing their dynamic activity states across cell types and tissues (together referred to as “cell types” for the rest of this work). Because each assay makes cell type-specific measurements, these assays must be performed for each cell type of interest separately. However, comprehensively profiling a large collection of cell types with assays targeting diverse attributes of chromatin is prohibitive due to practical constraints on material, cost and personnel. Hence, even the largest repositories of epigenomic and transcriptomic data are still incomplete in the sense that they are missing tens of thousands of potential experiments [1, 2, 3, 4, 5, 6].

To address this challenge, predictive models for imputing missing datasets have been proposed as an inexpensive and straightforward way to obtain complete draft epigenomes [7, 8, 9, 10, 11]. These models leverage the complex correlation structure of signal profiles from available experiments to impute signal for experiments that have not yet been performed. Recently, imputation models have been scaled to impute tens of thousands of experiments [12, 13] spanning dozens of assays in hundreds of human cell types. Although progress has clearly been made in developing imputation approaches, the field has thus far only explored a small portion of the space of potential imputation models. Notably, only one of the five methods surveyed above uses nucleotide sequence as input when making imputations.

We organized the ENCODE Imputation Challenge to encourage active development of imputation models. The challenge consisted of two stages and participants were encouraged to share ideas and reorganize into new teams between stages. In the first stage, participants were ranked based on their ability to impute a fixed validation set consisting of experiments randomly selected from within our data matrix. The second stage also measured imputation performance on a held-out set, but with two crucial differences from the first stage: first, the test data was collected during the challenge to ensure a truly prospective evaluation, and second, the test data was collected almost exclusively for poorly characterized cell types (only three of the 12 cell types in the test set have more than two training experiments).

Our initial expectation was that this challenge would primarily serve as an analysis of the components of imputation models and, ultimately, identify those that worked well. However, we found that fairly evaluating the imputations in the second stage was much more challenging than expected, and so the challenge instead served as an impetus to describe, and correct, distributional shifts in large collections of genomics data sets. Specifically, we found that a distributional shift occurs between the more recently collected paired-end data and the older single-end data available on the ENCODE portal due to small processing differences that have a big effect. Without correcting for this difference, we found that a baseline method outperformed all but two of the submissions using the performance measures defined before the challenge began, and those two submissions only performed marginally better than the baseline. After correction, more than half of the participants outperformed the same baseline.

We identified three key challenges in fairly evaluating imputation methods. First, differences over time in experimental procedure or data processing create distributional shifts across experiments which must be corrected for ensure a fair evaluation, and this correction must be more than a simple rescaling of the signal. This concern is particularly important when dealing with data sources, like the ENCODE Portal, that contain data collected over long periods of time. Second, while epigenomic imputation is most useful for cell types with few experiments, previous imputation work was evaluated using k-fold or leave-one-out cross-validation applied to an entire compendium. These evaluation settings over-emphasized the performance on well-characterized cell types and, unfortunately, good performance on well-characterized cell types is not always an indicator of performance on poorly characterized ones. Third, although designing several performance measures is necessary to capture the many aspects of a high-quality experimental readout, designing these measures without accounting for the first two issues can introduce redundancy in the measures, limiting their usefulness. We anticipate that giving proper consideration to these three issues in future works will be crucial for developing imputation methods that perform the best in practice.

Accordingly, this work focuses on characterizing the effect that these issues had on evaluating imputation methods, with the goal of providing guidance on how to fairly evaluate such methods in the future. When collecting a test set, one should ensure that processing steps have been uniformly applied to raw data and that the data have been collected using similar procedures. When differences in processing arise that cannot be undone, we propose handling distributional shifts by using a quantile normalization approach that separately normalizes signal in peaks and signal in background. We also propose a set of new performance measures that focus on orthogonal aspects of imputation performance. Finally, we note that performance that does not generalize from well characterized cell types to poorly characterized ones does not have a simple fix like the other issues do. Rather, this disparity can only be evaluated by explicitly including both well- and poorly-characterized cell types in the evaluation. At a higher level, one should ensure that at least one setting used to evaluate their approach matches how they expect the method to have the most impact in practice, namely, on poorly characterized cell types.

## 2 Methods

### 2.1 The ENCODE Imputation Challenge

We acquired candidate imputation models by hosting the ENCODE Imputation Challenge (https://www.synapse.org/encodeimpute), a public challenge for imputing epigenomic profiles, which began on February 20th, 2019, and concluded on August 14th, 2019. The challenge evaluated how well predictive models could impute held-out epigenomics experiments using other functional genomics experiments and nucleotide sequence as input (see challenge site for more details). Overall, we acquired 267 data sets from the ENCODE Portal to use as the training set, 45 data sets from the ENCODE Portal to use as a validation set, and performed 51 new experiments to use as a test set for prospective evaluation (Figure 1, Additional File 1).

**Figure 1:**
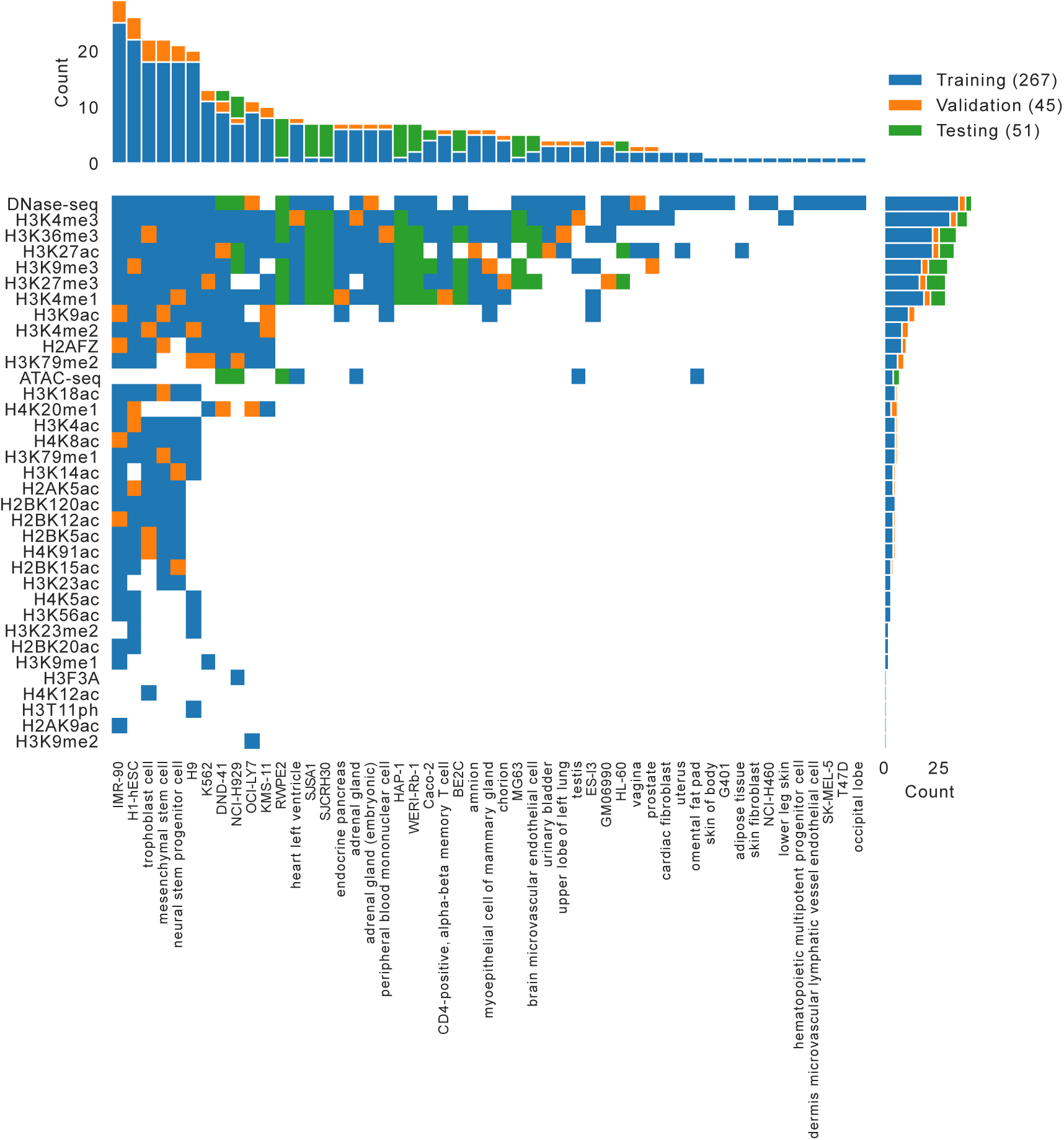
The challenge data matrix. The matrix shows the experiments used in the challenge, colored based on whether they were in the training set (blue), the validation set (orange), or the blind test set (green). White squares indicate that an experiment has not yet been performed. The marginal bar plots show the number of experiments in each assay and cell type.

The challenge was divided into two stages. In the first stage, participants were provided with the training and validation data sets as well as a real-time public leaderboard of performance on the held-out validation set. Team BrokenNodes and Hongyang Li and Yuanfang Guan tied for first place at the conclusion of the first stage (Additional File 2 Supplementary Figure S1, Additional File 3). In the second stage, the teams were allowed to re-organize, and participants were encouraged to refine their models using lessons learned from the first stage. The winners of the second stage, and of the entire challenge, were the top three teams based on performance on the held-out prospective test set, which the teams did not have access to.

The challenge was well attended with 196 people signing up on Synapse. Eight teams submitted results for the first round. After teams merged before the second round, 23 imputation models were submitted. Of these models, only one did not submit the full set of required imputations. Although our method for calculating team rankings as a part of the challenge accounted for missing imputations, our subsequent analyses excluded this model.

### 2.2 Performance Measures

Prior to the start of the challenge, we specified nine different performance measures to be used in the challenge. These performance measures included (1) the genome-wide mean-squared-error (MSE), (2) the genome-wide Pearson correlation, (3) the genome-wide Spear-man correlation, (4) the MSE calculated in promoter regions defined as ±2kb from the start of GENCODEv38 annotated genes [14], (5) the MSE calculated in gene bodies from GENCODEv38 annotated genes, (6) the MSE calculated in enhancer regions as defined by FANTOM5 annotated permissive enhancers [15], (7) the MSE weighted at each position by the variance of the experimental signal for that assay across the training set, (8) the MSE at the top 1% of genomic positions ranked by experimental signal, and (9) the MSE at the top 1% of genomic positions ranked by predicted signal. We note that 8 and 9 make a calculation similar to recall and precision, respectively.

We used a multi-stage process, originally developed for the ENCODE Transcription Factor Binding Challenge [16], to aggregate these performance measures into a single score to determine the challenge winners. First, ten equally-sized bootstraps were drawn from the pool of all genomic positions, and each of the nine performance measures was calculated for each team on each of the bootstraps for each experiment. For each bootstrap-experiment pair, the scores were converted to rankings across teams for each performance measure, and these rankings were then averaged across performance measures. This resulted in a score for each team in each bootstrap-experiment pair. This score was then converted back into a ranking over teams for each bootstrap-experiment pair. Next, these rankings were aggregated across experiments by calculating 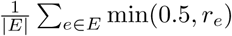 where *E* is the set of all experiments, *e* is an individual experiment, and *r*_*e*_ is a team’s ranking on experiment *e* divided by the number of teams. Finally, a rank was calculated across teams for each bootstrap, and the 90th percentile score, i.e. the second-best bootstrap rank, was used to determine the winners. This procedure is implemented at https://github.com/ENCODE-DCC/imputation challenge.

### 2.3 Baseline Methods

The methods submitted by the participants were compared to two baseline methods. The first baseline was the average activity, which is a straw-man imputation approach that simply predicts the average training set signal at each position in the genome across all cell types for a given assay type [17]. Consequently, this approach cannot make cell type-specific predictions. However, it represents the simple rule that regions of the genome that always exhibit peaks in signal and that regions of the genome that never exhibit peaks will continue to do so in other cell types. The second baseline was the Avocado model, using the same model architecture and training procedure described by Schreiber et al [12]. Importantly, this model was not tuned for this data set—it was applied as-is using the default settings and hyperparameters.

Although we had initially expected that ChromImpute [7] would serve as a baseline in this challenge, for logistical reasons ChromImpute was not applied to the challenge data until well after the challenge concluded. Because the participants did not have access to these predictions, as they did with the other two baselines, we did not include ChromImpute in the original rankings or analysis. However, we have included a ranking of methods that includes ChromImpute in a re-analysis of the challenge participants using six of the measures used to evaluate the original ChromImpute method [7], as a reference (Additional File 2 Supplementary Figure S2/S3). These measures emphasize the relative distribution of signals, and included Pearson correlation, three measures quantifying percentage overlap between positions exhibiting high signal, and AUC measures for predicting peaks in observed signal from imputed signal values and vice versa. In order to obtain team ranks on these measures, we first ranked each team’s prediction for each test track on each measure separately. We then averaged ranks across metrics and re-assigned integer ranks in each track for each team. Each team’s final rank was then computed from the average of their predictions’ track ranks for the 51 test tracks.

### 2.4 Quantile Normalization

We developed a three-step quantile normalization method for normalizing signal across genomics experiments. Because signal distributions differ significantly across assays, we applied this normalization separately for each assay. Importantly, the normalization is also done separately for signal in peak and background regions (as defined by MACSv2 peak calls for the experiment [18]) to account for peaks spanning differing proportions of the genome across cell types. In the first step, quantiles are derived separately from each training set experiment. That is, if there are *N* training set experiments, *M*_*p*_ peak quantile bins, and *M*_*b*_ background quantile bins, one would extract 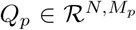 and 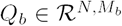. Quantiles are extracted by ranking all signal values for an experiment (in peaks or outside of peaks, respectively), binning those ranks into either *M*_*p*_ or *M*_*b*_ equally sized bins, and assigning to each bin the average signal value from positions within the bin. In the second step, an average is taken across experiments for each quantile bin to construct reference quantiles 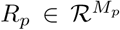 and 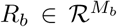. Finally, *R*_*p*_ and *R*_*b*_ are applied to the test set tracks, with *R*_*p*_ being applied only within signal peaks and *R*_*b*_ being applied only within background regions. Because peak regions are more complex and span a larger dynamic range than the background, we set *M*_*p*_ to be 1000 and *M*_*b*_ to be 50. Given that this procedure is designed to combat distributional shift, we note that it should be applied to test set experiments before evaluation.

### 2.5 Data Processing

We processed the DNase and ATAC-seq experiments using a uniform pipeline [19]. First, FASTQ files containing read sequences and quality scores for the training and validation sets experiments were downloaded from the ENCODE Portal, and FASTQs for the test set experiments were acquired from our own experiments. For ATAC-seq experiments (but not DNase-seq), we first trimmed adapters and then mapped reads to the hg38 reference human genome using the Bowtie2 [20] aligner. After mapping, reads were filtered to remove unmapped reads and mates, non-primary alignments, reads failing platform/vendor quality checks, and PCR/optimal duplicates (-F 1804). Reads mapping reliably to more than one location (MAPQ *<* 30), i.e., multi-mapping reads, were removed. Duplicate reads were then marked with Picard MarkDuplicates [21] and removed. For single-end DNase data sets, a single read was chosen from a set of duplicate reads, whereas for paired-end data sets, read-pairs were chosen if any one of the two reads in the pair was unique. Although this is the standard approach for de-duplicating single-end and paired-end data, this step had unintended consequences for the challenge, which we describe in Section 3.2. For ATAC-seq data, 5’ ends of filtered reads on the + and - strand were shifted by +4 and -5 bp respectively to account for the Tn5 shift. Reads from biological and technical replicates were merged. We normalized the sequencing depth across data sets by subsampling them to a maximum of 50 million reads (after excluding reads mapping to mitochrondria). Although there are several ways to represent the signal from sequencing experiments, e.g., read-counts and fold-change, we chose to use the statistical significance of the fold-change to be consistent with previous imputation literature [11, 12, 7, 10]. We used the MACSv2 peak caller to compute the fold-enrichment and statistical significance. MACsv2 was applied to smoothed counts (150 bp smoothing window) of read-starts (5’ ends of reads) at each position in the genome relative to the expected number of reads from a local Poisson-simulated background distribution. We filtered out all peaks that overlapped with the ENCODE Exclusion list consisting of abnormal high signal regions [22]. We provided the genome-wide signal tracks containing the statistical significance of enrichment (i.e., the -log10 p-values) at each basepair in the genome. The processing pipeline is open-source and available at https://github.com/ENCODE-DCC/atac-seq-pipeline.

Next, we processed the histone ChIP-seq experiments using the ENCODE processing pipeline [23]. For each experiment we downloaded FASTQ files from the ENCODE Portal for at least two replicate experiments and a control experiment. All reads were mapped to the hg38 reference human genome using the BWA aligner [24]. After mapping, the process was similar to the ATAC-seq/DNAse-seq pipeline. Reads were filtered to remove unmapped reads and mates, non-primary alignments, reads failing platform/vendor quality checks, and PCR/optical duplicates (-F 1804). Multi-mapping reads (MAPQ *<* 30) were also removed. Duplicates were identified using Picard MarkDuplicates and subsequently removed, with the same single-end vs. paired-end differences as mentioned for DNase data sets. Reads from the biological and technical replicates were then merged. We normalized the sequencing depth across data sets by subsampling each to a maximum of 50 million reads. We used the MACSv2 peak caller to calculate fold-enrichment and statistical significance of counts of extended ChIP-seq reads (reads were extended in the 5’ to 3’ direction based on the predominant fragment length), relative to the number of extended reads from the control experiment, and filtered out peaks that overlapped with the ENCODE Blacklist [22]. The statistical significance of the enrichment was computed using a local Poisson null distribution whose mean parameter is estimated from the control experiment. For the purposes of this challenge, we provided the genome-wide signal tracks containing the statistical significance of enrichment (i.e., the -log10 p-values) at each basepair in the genome. The processing pipeline is open-source and available at https://github.com/ENCODE-DCC/chip-seq-pipeline2.

## 3 Results

### 3.1 The ENCODE Imputation Challenge

Participants submitted 23 models to the second stage of the challenge (see Section 2.1). Each group was allowed to submit up to three models to encourage inclusion of unorthodox solutions with at least one submission. As a result, the models encompassed a diverse range of strategies (see Table 1). The models differed primarily along three axes. The first axis was the signal preprocessing, with almost every method further preprocessing the data from the given -log10 signal p-values. The second axis was the data sources used to construct input features. Most methods followed previously published methods by only using assay measurements as inputs (denoted “functional” in Table 1). However, five of the methods used nucleotide sequence as input, eight methods used the average activity baseline, and three used Avocado’s imputations. The third axis was the manner in which the underlying tensor structure of the data was modeled. Some methods explicitly modeled the data as a tensor (e.g., imp and Lavawizard), whereas other methods only implicitly modeled the structure through rule-based approaches or similarity methods (e.g., the HLYG and KNN-based approaches).

**Table 1:** Methodologies of imputation methods. The table lists the modeling strategies and input features used by each of the models, as reported by the teams. The models include k-nearest neighbors (KNN), deep tensor factorization (DTF), autoencoders (AE), convolutional neural networks (CNN), hidden Markov models (HMM), and gradient-boosted decision trees (GBT). The authors of Aug2019Impute and CostaLab v2 did not describe their methods.

| Name | Model | Norm | Sequence | Inputs |  |  |
| --- | --- | --- | --- | --- | --- | --- |
|  |  |  |  | Functional | Average | Avocado |
| Aug2019Impute |  |  |  |  |  |  |
| BrokenNodes/v2 | KNN | arcsinh |  | ✓ |  |  |
| BrokenNodes v3 | KNN | arcsinh |  | ✓ |  | ✓ |
| CostaLab v2 |  |  |  |  |  |  |
| CUImpute1/CUWA/ICU | ensemble | arcsinh |  | ✓ | ✓ | ✓ |
| Guacamole/Lavawizard | DTF | arcsinh |  | ✓ | ✓ |  |
| HLYG/v1/v2 | GBT | quantile | ✓ | ✓ | ✓ |  |
| imp/imp1 | DTF+AE | Cauchy |  | ✓ |  |  |
| KKT-ENCODE | CNN | arcsinh | ✓ |  |  |  |
| LiPingChun | DTF | arcsinh |  | ✓ | ✓ |  |
| NittanyLions | KNN |  |  | ✓ |  |  |
| NittanyLions2 | KNN | quantile |  | ✓ |  |  |
| SongLab | CNN | log1p |  | ✓ |  |  |
| SongLab2 | HMM |  |  | ✓ |  |  |
| SongLab3 | CNN | log1p |  | ✓ | ✓ | ✓ |
| UIOWA | CNN | quantile | ✓ | ✓ |  |  |

An initial inspection of the imputations revealed that most methods captured the general shape of the signal well. Examples drawn from H3K27ac in brain microvascular endothelial cells and DNase-seq in DND-41 cells (Figure 2A/B, Additional File 2 Supplementary Figure S4) suggest two sources of error: the misprediction of a small number of peaks relative to the total number of true peaks, and the misprediction of the precise signal value within correctly predicted peaks. Focusing on the misprediction of peaks, we noted that some methods made similar mistakes as the average activity baseline, whereas others made similar mistakes as the Avocado baseline (gray highlights in Figure 2A/B). Unsurprisingly, methods that used Avocado’s imputations as input had the highest genome-wide correlation with Avocado’s predictions (it is worth noting that CUImpute1 only used Avocado’s imputations for some, but not all, assays). In contrast, methods that explicitly used the average activity did not always exhibit higher correlation with it than other methods (Supplementary Figure S5). This finding suggests that, because the average activity can be directly derived from the training set, many types of models are able to implicitly learn it even when not explicitly trained on it.

**Figure 2:**
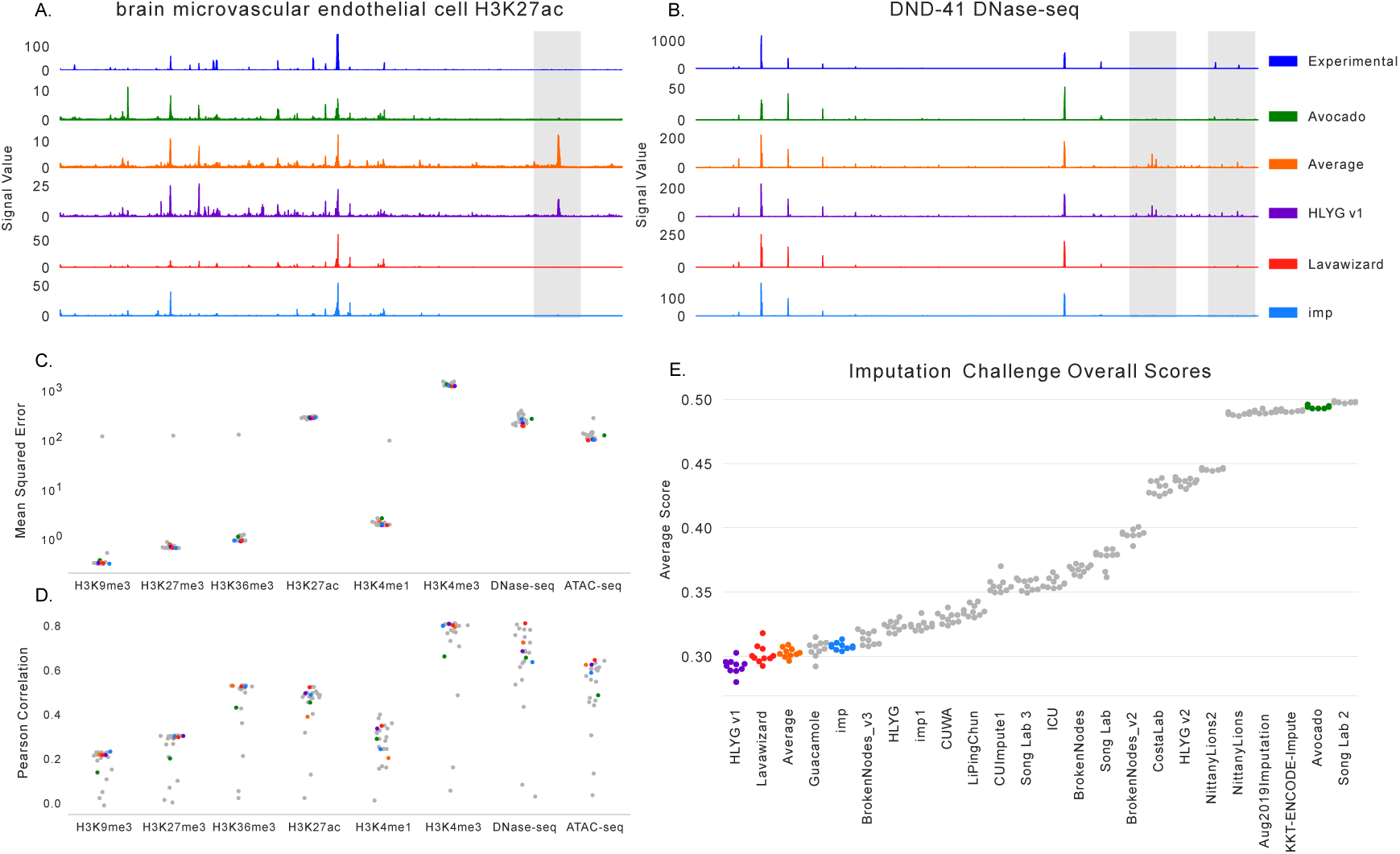
Results from the ENCODE Imputation Challenge. (A) The H3K27ac signal for brain microvascular endothelial cells that is observed (in blue), from baseline methods, and from the winning three teams in the challenge. (B) The same as (A) except for DNase-seq signal in DND-41 cells. (C) The average MSE for each method across test set tracks and bootstraps but partitioned by assay type. (D) The same as (C) except for Pearson correlation. (E) The overall score, calculated as described in Section 2.2, across all test set tracks and performance measures shown for each bootstrap for each team. The baseline methods and winners are colored.

Next, we comprehensively evaluated the methods using a battery of performance measures that were specified at the beginning of the challenge (see Section 2.2, Additional File 4). We found that performance on these measures depended heavily on the imputed assay (Figure 2C/D). For instance, most models exhibited four orders of magnitude higher MSE on H3K4me3 than on H3K9me3. However, several assays that exhibited the highest MSE also exhibited the highest Pearson correlation, indicating that the scale of MSE across assays is likely more related to the dynamic range of the assay rather than the accuracy of the imputations. Unsurprisingly, a projection of all imputed and experimental tracks clustered predominately by assay type (Silhouette Score = 0.4601), as opposed to by cell type (SS = -0.4028) or imputation method (SS = -0.3133, Supplementary Figure S6). Accordingly, we used a rank-based transform to account for differences in dynamic range when calculating global performance measures across experiments (see Section 2.2) to ensure that assays with large dynamic ranges did not dominate the evaluation. After calculating the global performance of each method, we found that there was a gradient of methods that performed increasingly well, and a set of methods that performed relatively poorly (Figure 2E). The best performing methods, and hence the winners of the challenge, were Hongyang Li and Yuangfang Guan v1 (abbreviated as “HLYGv1”) in first place, Lavawizard and Guacamole (two similar methods from the same team) tied for second place, and imp in third place.

Given the diverse modeling strategies of the winning teams, our primary take-away from these results is that there does not appear to be a single key insight that led to good overall performance on the measures used in the challenge. HLYGv1 used nucleotide sequence as input, but so did KKT-ENCODE and UIOWA Michaelson; all three models submitted by Hongyang Li and Yuanfang Guan used gradient boosted trees (GBTs), yet their models exhibited both good and poor performance. However, these results do suggest certain models to be wary of: convolutional neural networks and k-nearest neighbor models underperformed deep tensor factorization (DTF) and GBT models. This is likely because the similarities used by KNN models are a less sophisticated version of the representations learned by tensor factorization approaches, and that the specific structure presented in the data is not well modeled by simple applications of convolutions.

However, when we compared model performance to the baseline methods, we made two important observations. First, almost every team outperformed the Avocado baseline, as one might expect because the participants had access to the Avocado model and predictions during the development process, and because the default settings were used for Avocado despite them being tuned for significantly larger amounts of training data. Second, the average activity baseline performed extremely well, coming in third in our ranking and first place in five of the nine performance measures used (Additional File 4). Both of these observations are a reversal from the first round in the challenge, where Avocado outperformed all the participants but almost all the participants outperformed the average activity baseline (Supplementary Figure S1). This reversal in performance between the two baselines is partially because the evaluation setting changed from overrepresenting well characterized cell types to focusing on poorly characterized ones and, as we will see later, partially due to the performance measures used for the challenge.

### 3.2 Accounting for distributional shift

A visual inspection of the test set experiments revealed significant distributional differences in peak signal values between the training and test sets for some assays (Figure 3A). Most obviously, the signal values within H3K4me3 peaks from test set experiments were generally much higher than the signal values within peaks from training and validation set experiments (Figure 3B). Although one would expect a locus to exhibit different signal in different cell types because of real biology, one would also expect that the distribution of signal values within peaks across entire experiments would be similar for experiments of the same assay. Because distributional shifts have major ramifications for the scale-based performance measures used in the challenge, we next investigated the source of these distributional differences.

**Figure 3:**
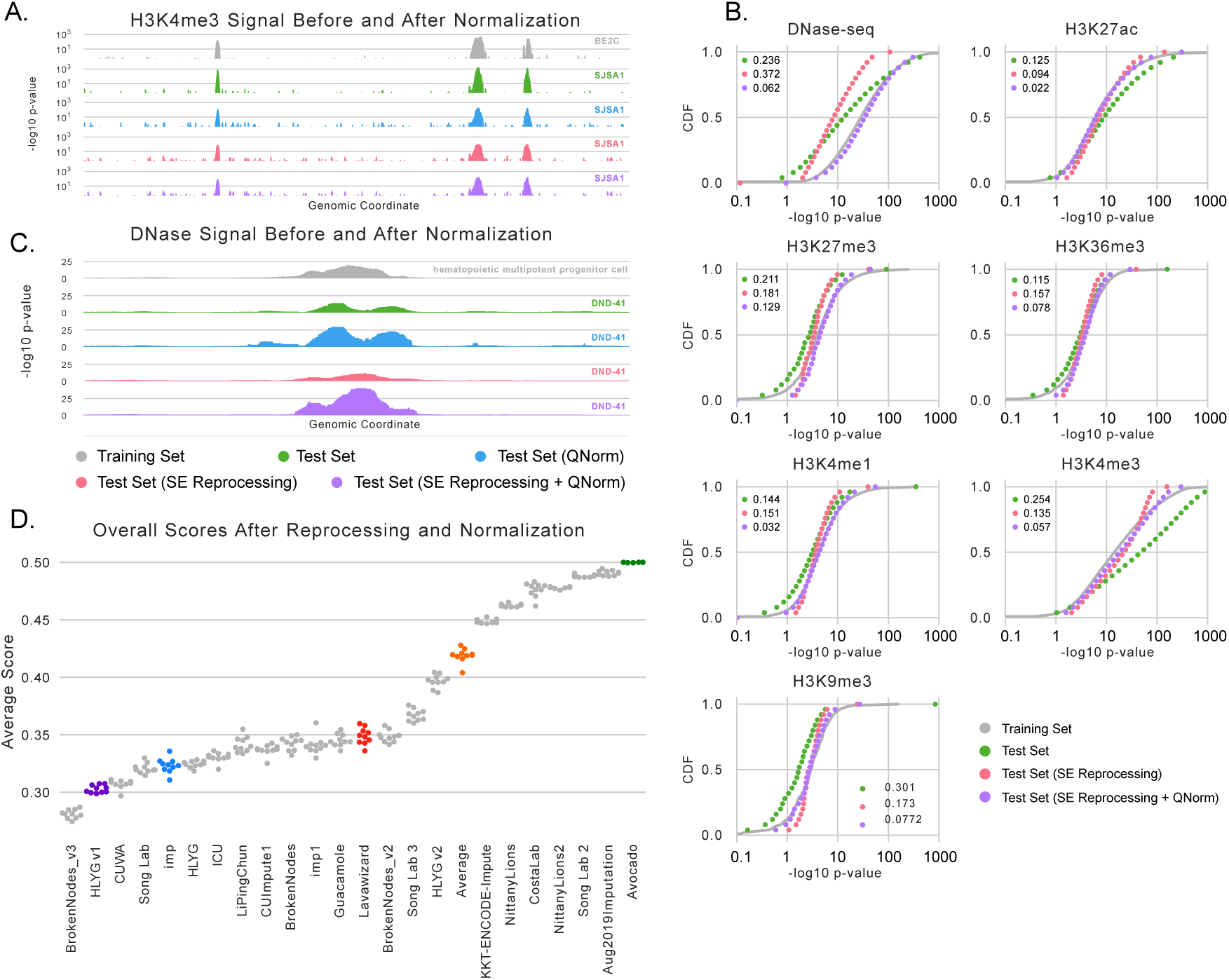
Distributional shift and quantile normalization. (A) Experimental signal measuring H3K4me3 in BE2C cells from an unnormalized training set experiment (gray), an unnormalized test set experiment in SJSA1 cells (green), the test set signal after quantile normalization (blue), the test set signal after single-end reprocessing (red), and the test set signal after single-end reprocessing and quantile normalization (purple). (B) Distributions of signal values within peaks in chr16/17 for each reprocessed assay across the unnormalized training set (gray), the unnormalized test set (green), the single-end reprocessed test set (red), and the single-end reprocessed and quantile-normalized test set (purple). The KS statistics between the training set distribution and the test set distributions are shown in the legends and the CDFs are summarized using 25 dots for visualization purposes. (C) An example locus that exhibits a DNase peak in both the training and test sets. (D) A rescoring of the challenge participants against single-end reprocessed and quantile-normalized test set signal.

After considering several potential covariates that could explain this distribution shift, including multiple measures of experimental quality (Additional File 2, Supplementary Figure S7), we found that the primary driver was a subtle difference in how the test set experiments were processed. By design, the test set experiments were performed during the challenge to ensure a truly prospective evaluation. However, experimental methods have changed in the many years since the training data were collected. Most notably, collecting paired-end data is now the standard approach for ENCODE data sets because the procedure yields higher quality data and is now cheap enough for broad usage; however, almost all of the training set experiments predate this switch and involve single-end data. The processing of single-end and paired-data data is largely similar, but a crucial difference occurs in the deduplication step. Specifically, deduplication of single-end reads using PICARD [21] allows the mapping of only one read start to each position on the genome on each strand. In contrast, deduplication of paired-end data can result in more than one read-start per position on each strand because read-pairs are only removed if the read start of *both* ends are duplicates. Consequently, the number of reads mapping within peaks from paired-end data can be significantly higher than what one would get using single-end data. Importantly, the shift is not simply caused by paired-end data being higher quality, as we first explored, but rather differences in the deduplication step.

We confirmed that differences in processing, rather than differences in data quality, explained the distributional shift by reprocessing the paired-end data sets (except for ATAC-seq which requires paired-end data) as single-end data. Specifically, for each paired-end experiment in the test set we concatenated the FASTQ files of reads from both ends and ran the same single-end processing pipeline that was run on the other single-end experiments in the challenge. We found that the reprocessed data had distributions of peak signal values significantly closer to the training set, as measured by the Kolmogorov-Smirnov (KS) statistic, for four of the histone modification assays including H3K4me3 (Figure 3B). The remaining two histone modification assays already resembled the training set before reprocessing. However, we found that the distribution of DNase-seq peak signal values had a larger KS-statistic after reprocessing than before. This is likely because 21 of the 38 training set experiments contained paired-end data, which would shift the distribution of signal values in the training set up. Although the most principled next step would be to reprocess all of the experiments used in the challenge and subsequently re-training and reevaluating each submission, this analysis was not possible because we only required that the three challenge winners submit code that could retrain their models on new data sets. Given no perfect solution, we chose to continue with the single-end reprocessed test set tracks for our subsequent analyses.

We found that reprocessing the histone modification data significantly reduced the distributional shift but did not perfectly correct it. The remaining differences are likely related to small changes in experimental protocol over time, such as improvements in sequencing technology, antibodies used, and read lengths measured. A general-purpose correction for the remaining differences is to explicitly quantile normalize the data such that the signal values in the testing experiments exhibit the same signal distribution as those in the training experiments. Quantile normalization is powerful because it is a non-linear method, in contrast to min-max or z-score scaling, and has been extensively applied to genomics data sets, including those measuring bulk gene expression [25], single-cell RNA-seq [26], and ChIP-seq data when combined with a spike-in reference [27]. We account for differing proportions of the genome exhibiting peaks across cell types by separately quantile normalizing the signal within peaks and the signal in background regions (see Section 2.4 for details). Finally, because the distribution of signal is significantly different across assays, we apply this quantile normalization to each assay separately. After normalization, we confirmed that the distribution of within-peak test signal values was almost identical to the distribution of within-peak training signal values across all assays (Figure 3B), even for the DNase-seq experiments.

In theory, one could apply quantile normalization to the original paired-end test set data and, by definition, produce signal values with the same distribution without the need for reprocessing. However, when looking at a representative DNase peak, we found that the reprocessed data was not a simple monotonic transform of the original data (Figure 3C). Specifically, the paired-end data exhibited a peak shape unlike that observed in the single-end data, and simply quantile normalizing the signal does not fix the differences in shape. More comprehensively, when considering a 10Mbp region of chr1 on each of the 48 reprocessed experiments, we clearly observed that paired-end data is not a monotonic transformation of single-end data (Additional File 2, Supplementary Figure S8). Although the assays associated with activity, such as H3K4me3 and DNase-seq, exhibit Spearman correlations up to 0.938 between the paired-end and single-end processed signals, repressive marks exhibit Spearman correlations as low as 0.037, and the average Spearman correlation across all tracks was only 0.453. Further, even though some assays exhibit high correlation, this value is inflated by the large number of low-signal values and, indeed, the largest variability comes at loci with high signal values.

Moving forward with our method of reprocessing the test data using single-end settings and then quantile normalizing to correct the remaining differences, we next re-scored the originally submitted imputations (Figure 3D, Additional File 5, Additional File 6). We observed that the number of methods outperforming the average activity baseline increased from two to 16 and that BrokenNodes v3 rose from sixth place to first place in the rankings. Although HLYGv1 remains within the top three, the other two winners descended in the rankings. This might be explained by HLYGv1 using quantile normalization, albeit a slightly different version than the one we used, during training. Interestingly, many of the methods performed similarly to each other, reinforcing the idea found in the original challenge that there is not necessarily one way to do imputation. Indeed, the best performing model is a simple KNN-based approach using arcsinh-transformed data and the second best performing model uses gradient-boosting trees on quantile transformed data. Critically, we note that it would not be fair to use these rankings to declare challenge winners because we did not give the teams an opportunity to retrain or tune their methods on the transformed data. Rather, our take-away is that the distributional shift is partially responsible for the good performance of the average activity baseline but does not fully explain it.

### 3.3 Designing more informative performance measures

Although the measures used in the challenge were devised to rank methods independently for each experiment based on their genome-wide (or across large portions of the genome) performance, this property meant that they ultimately exhibited a high degree of redundancy with each other (Supplementary Figure S9). Essentially, by uniformly weighting all positions along the genome, methods with low genome-wide MSE were likely to have low MSE within promoters, gene bodies, or the top 1% of signal as well. Exacerbating this issue, MSE-based measures were disproprotionately confounded by the large distributional shift described in the previous section in comparison to the shape-based measures. Illustrating this, we found that most of the residual—sometimes over 99% in H3K4me3 assays—came at correctly-predicted peaks (Supplementary Figure S10A). Realizing this weakness, we next designed three new types of performance measures that, respectively, reweighted genomic bins based on signal strength, considered multiple experiments simultaneously, and focused on shape within active areas. All evaluations in this section are done against the reprocessed, quantile-normalized test set signal.

#### 3.3.1 Partitioning by signal strength

A strategy for measuring performance in a complementary way to uniformly-weighted genome-wide performance is to explicitly calculate the performance with respect to the magnitude of either the observed or imputed signal (Figure 4A/B). Rather than being limited by considering only the top 1% bin of signal, such as by using the mse1obs or mse1imp measures, considering all signal bins provides a finer-grained view of model performance. As an example, if the imputations exhibit high accuracy when the imputed signal is high, then one may be confident that predicted peaks are correct when using imputations for which there is no corresponding experimental data; in contrast, if the imputations exhibit low accuracy when the imputed signal is high but higher accuracy when the imputed signal is low, then one might be more skeptical of imputed peak calls but more trusting of regions not called as peaks, e.g. facultative peaks that are not active in the studied cell types. Although any measure can be partitioned by signal magnitude, we focus on accuracy between binarized imputed signal and peak calls for the experimental signal. Accuracy was excluded from the original set of performance measures because the sparsity of peaks can make it difficult to interpret genome-wide; in this setting, we anticipate accuracy to be more valuable in the signal bins where one might reasonably find a peak. Importantly, we did not use rank-based classification measures (e.g., AUROC or AUPR) here, because once the signal is partitioned by strength, applying a rank-based measure to each bin is less meaningful than when applied genome-wide.

**Figure 4:**
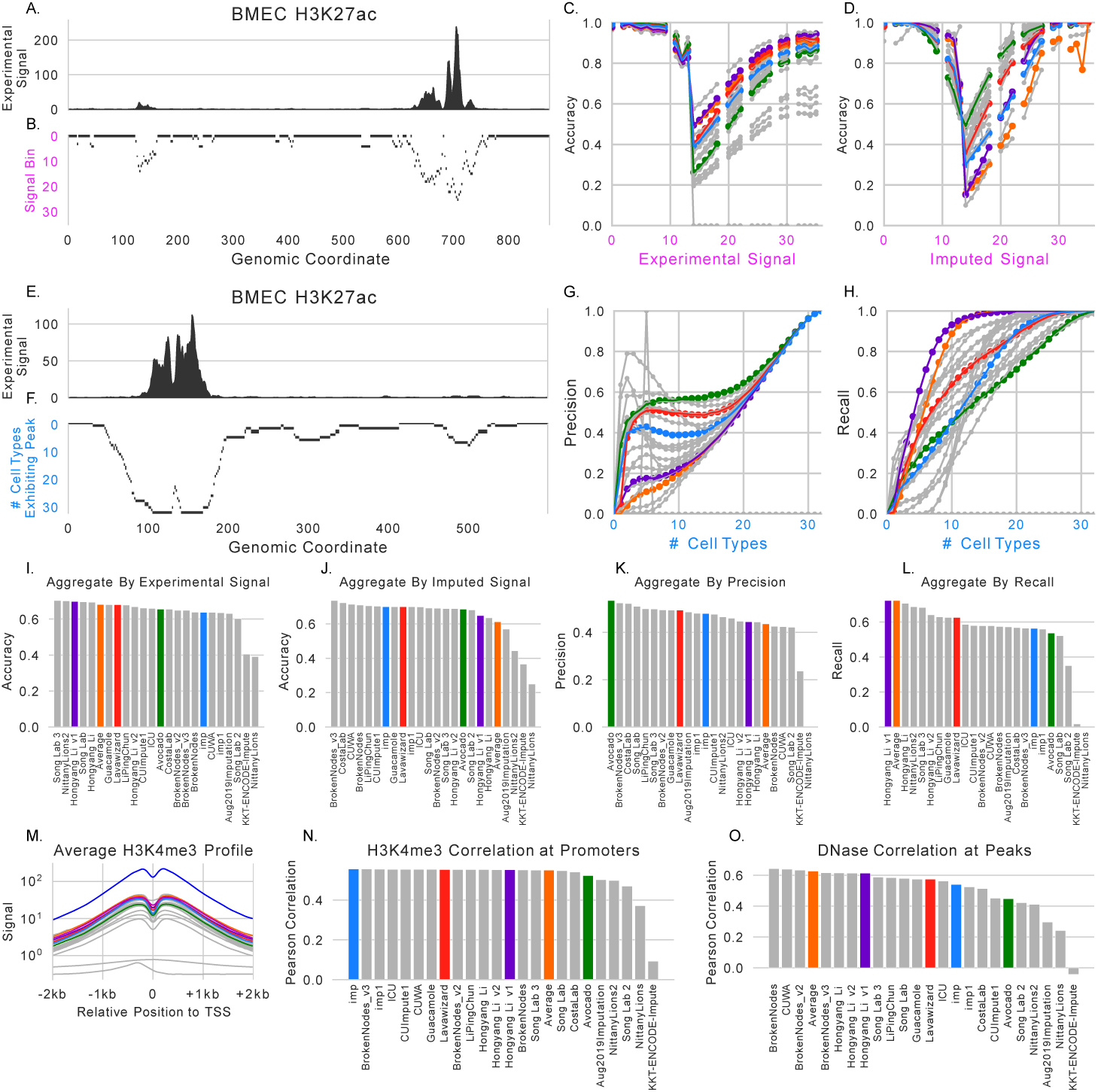
Additional performance measures. (A) Experimentally observed signal for H3K27ac in brain microvascular endothelial cells. (B) An example of partitioning the track from A into logarithmically-spaced bins (the rows). (C) The accuracy between binarized imputations and MACS2 peak calls for each signal bin when using the experimental signal to define the bins. (D) The same as C except using the imputed signal to define the bins. (E) The same as A but a different locus. (F) The same as B except calculating bins using the number of cell types that each locus exhibits a peak in. (G) The precision of the binarized imputed signal against MACS2 peak calls when evaluated separately for each bin. (H) The same as H except the recall instead of the precision. (I) The average area under the curves, calculated as shown in C, across all test set tracks for each participant. (J) The average area under the curves calculated as shown in D across all test set tracks for each participant. (K) The precision score calculated in the same manner as I/J. (L) The same as K, except the recall score. (M) The average H3K4me3 profile of experimental (blue), quantile-normalized (magenta), and imputed signals at strand-corrected promoters. (N) The average Pearson correlation between imputed and quantile-normalized signal across all promoters and H3K4me3 test set tracks. (O) The average Pearson correlation between imputed and quantile-normalized signal across all observed DNase peaks and DNase test set tracks.

When we partitioned genomic loci based on experimental signal, we found that model performance aggregated across all tracks generally falls into three regimes: (1) when the imputed or experimental signal is low, the accuracy is high, (2) when the imputed or experimental signal is between 1 and 10 the accuracy severely drops, and (3) when the imputed or experimental signal is high, the accuracy returns to being high (Figure 4C). Although the second regime includes ambiguous peak calls, it also includes the most difficult to call peaks (thus, the relatively low accuracy) and should be emphasized by performance measures. When focusing on H3K27ac signal in brain microvascular endothelial cells we can also see that rankings flip between the first and second regime (Figures 4C-D); imp1 and LavaWizard both outperform the average activity and HLYGv1 when the experimental signal is low, but perform significantly lower when the experimental signal is higher.

Interestingly, the ranking of methods is almost reversed when partitioning genomic loci using the imputed signal instead of the experimental signal (Figure 4D). HLYGv1 and the average activity are among the top performers when partitioning by experimental signal but are among the worst performers when partitioning by imputed signal. An explanation for this flip is that these approaches measure notions of precision and recall, respectively, which have a known trade-off. Because the average activity is essentially a union of peaks across cell types in the training set, it will have a high recall but a low precision. Methods, such as HLYGv1, that rely too heavily on the average activity will exhibit the same tradeoff (Figure 4C/D).

A straightforward way to condense these curves into a single value for a performance measure is to take the average value across the curve. This value is essentially a re-weighting of genome-wide accuracy that uniformly values each bin of signal values rather than each locus, and so will downweight the more common low signal value loci and upweight the less common higher signal values ones. Notably, the winners of the ENCODE Imputation Challenge did not perform the best across all test set experiments when partitioning by either experimental signal or by imputed signal (Figure 4I/J). Indeed, the top two performers when partitioning by experimental signal (Song Lab 3 and NittanyLions2) came in 12th and 19th respectively in the original evaluation.

#### 3.3.2 Prediction of facultative peaks

A primary source of error for imputation models comes from loci that exhibit functional activity in some, but not all, cell types. Evaluating whether the imputations can distinguish between cell types that do and do not exhibit signal at a given locus is crucial for ensuring that the imputations are cell type-specific. However, because traditional genome-wide performance measures treat each experiment independently, they cannot explicitly evaluate this property. To better understand how well these methods can identify what cell types loci are active in, for each assay we partitioned genomic positions by the number of experiments that exhibit a peak for that assay and then evaluated each partition separately. For example, if a locus exhibited a DNase-seq peak in 3 out of 5 cell types, that locus would be grouped for evaluation with other loci that also exhibited DNase-seq peaks in 3 out of 5 cell types (Figure 4E/F). This analysis is similar to the one presented by Schreiber et al. [11]

We observe trends that are reminescent of partitioning loci by signal strength. As the number of cell types that exhibit peaks increases, so too does the precision and recall of the methods (Figure 4G/H). This indicates that, generally, imputation methods are better at predicting peaks at facultative peaks than they are at predicting cell type-specific activity. Interestingly, we noted that several methods had peaks in precision when the number of cell types the peak was expressed in is low. Given that performance was extremely variable in this regime, we think that focusing on this measure in future studies will be useful when comparing models. Consistent with the role that the average activity plays as essentially the union of peaks across cell types, we see that it has a low aggregate precision score across all test set tracks but has the second highest aggregate recall score (Figure 4K/L). Put another way, the average activity is very good at identifying peaks that are common across many cell types but very poor at identifying the cell types that cell type-specific peaks occur in. Somewhat surprisingly, the Avocado baseline had the highest aggregate precision score, but the challenge winners that most resemble it (Lavawizard and imp) did not exhibit the most similar performance.

#### 3.3.3 Relative peak shape

The performance measures that have been proposed so far predominantly involve genome-wide calculations, even if they involve re-weighting loci contributions. An alternate form of performance measure is to focus on specific forms of biochemical activity at loci that are known to be relevant. The MSEProm, MSEEnh, and MSEGene measures attempt to quantify this by focusing on promoters, enhancers, and gene bodies respectively, but measure the performance of all assays at these loci. Next, we investigate two more performance measures that follow the reasoning of Ernst et al. [7] that only specific assays should be measured at these loci.

The first measure evaluates the shape of H3K4me3 signal at promoter regions. This histone modification is known to be enriched at promoter elements and is indicative of active transcription. Further, after correcting for the strand of the promoter, the mark exhibits a distinctive bimodal pattern (Figure 4M). We reasoned that focusing on the ability to recapture this shape would provide an orthogonal evaluation to the other performance measures proposed so far. We calculated the average Pearson correlation between the imputed signal and the quantile-normalized experimental signal across all gene promoters for all test set tracks measuring H3K4me3. Most of the methods outperformed the average activity baseline but only one of the challenge winners were in the top five according to this measure (Figure 4N).

The second measure evaluates the shape of DNase signal at observed DNase peaks. We anticipated that recapturing the shape of DNase signal would be more challenging because DNase does not exhibit a pattern that is as consistent as H3K4me3 at promoter regions. Further, the subtle patterns encoded in DNase signal can be useful for deciphering the precise regulatory role that the underlying nucleotide sequence is playing. Consistent with predicting DNase signal being a more challenging task, we found that methods exhibited a wider range of performances than they did with H3K4me3 prediction (Figure 4O). We also found that only three methods outperformed the average activity baseline. This might initially be counterintuitive, because chromatin accessibility is fairly cell type-specific. However, because this evaluation is limited to observed DNase peaks, methods are not being penalized for incorrectly predicting that non-peak regions are exhibiting peaks. This observation indicates that accessible loci largely retain the shape of their peaks across cell types when binned at 25 bp resolution.

## 4 Discussion

A central theme of this work is that evaluating models that rely on large collections of genomic data sets can be more difficult than one might initially expect and, consequently, that results can be confounded even when one does not make any obvious mistakes. In our analysis, we identified three issues that made analysis of imputation models more difficult than we initially thought: distributional differences in the underlying data, previous evaluation focusing on well-characterized cell types and in larger compendia, and performance measures that were either redundant or sensitive to the first two issues. We addressed these issues by proposing a quantile normalization approach that treats peak and background signal separately, and proposing new performance measures that were less redundant with each other and covered more aspects of performance than the original measures.

When the challenge was originally designed, the participants were not required to submit working code in order to lower the barrier to entry and allow participants to use their own custom hardware. Although this likely increased participation, it also caused a recurring problem in our later analyses because we could not retrain models on reprocessed data, or on different subsets of data. For example, reprocessing all the data using the single-end settings would likely have been the correct thing to do from a theoretical point of view, but was impossible as a practical matter because we did not have the required code. Likewise, we had hypothesized that part of the reason for changes in rankings between the first and second stages (including in our baselines) was because the first stage involved evaluation on a randomly selected held-out test set of experiments, which are biased towards well-characterized cell types, and the second stage explicitly evaluated only poorly characterized cell types. Because we could not re-train the models and evaluate them on cell types giving variable amounts of information, we could not comprehensively pursue this line of inquiry using the challenge data.

Based on our experience running this challenge, we have several recommendations for the organizers of future challenges involving genomic data sets. First, ensure that participants are compared against naive baselines such as the average activity. Without this baseline, we might not have identified as easily the distributional shift or the worse performance on sparsely characterized cell types. Second, participants should be required to submit code that can reproduce the training of their models so that more in-depth analysis can be done later. Potentially, the organizers should provide a scaffold that the participants fill in with their own code so that the organizers do not need to decipher each submission to use it properly. Third, organizers should explicitly look for distributional shifts across data splits, and even between pairs of data sets, as a quality control step. For example, paired end datasets from cancer cell lines can often contain large regional distribution shifts and outliers driven by cell line-specific copy number variation. Even when these shifts are explained by biological processes rather than experimental biases, tailoring an analysis that accounts for these shifts can be an important aspect of a fair evaluation. Finally, organizers should design performance measures that have minimal redundancy with each other, potentially as measured using the average activity before the challenge begins. Naturally, without a singular end-goal in mind it can be difficult to balance the various aspects of performance in a manner that will satisfy everyone, but having redundant performance measures is clearly not helpful.

An unaddressed, but important, issue is determining the most informative target for imputation methods to predict. The most common target in imputation literature has been the statistical significance from a peak-calling algorithm. Predicting the statistical significance can be more informative than predicting read counts directly because read counts can suffer from unwanted experimental biases and the peak-calling algorithm can explicitly consider a control track. Our challenge setting is consistent with that literature. However, an issue with predicting p-values is that fewer tools take those as input than take read counts as input. In fact, performing peak calling using imputations is not obvious because it is unclear that simply thresholding the uncalibrated p-values is the correct approach. Potentially, future iterations of the imputation work could involve imputing read counts but allowing models to directly incorporate the control tracks and other covariates such as sequencing depth, single-end or paired-end status and data quality metrics as well [28]. Although there would be some engineering challenges with such a task, such as designing alternate loss functions or performance measures based on counts, imputation of read counts might be more readily adopted.

Although the issues we described made the analysis of the results of this challenge more difficult, we made several important findings that we hope will guide the design and analysis of predictive models that rely on genomics data in the future. Specifically, even outside the context of a challenge, being aware of distributional shifts and evaluating a newly proposed model with a wide set of performance measures can help ensure that the model is robust in practice. Further, the difficulties that we faced are not unique to the setting of imputation. Indeed, these issues can affect any model that is trained or evaluated using large collections of publicly available data sets.

## 4.1 Author Contributions

A.K., C.B., J.S.S., M.K. and W.S.N. designed the challenge. J.L. processed the data for the challenge. J.L. and J.S.S. provided technical support for the challenge. A.K., W.S.N., J.S., and C.B. ran the challenge. J.S. and C.B. manually validated the challenge winners. J.S. designed and performed the subsequent analyses after the challenge concluded and wrote the manuscript. C.B., A.K. and W.S.N. edited the manuscript.

H.L. and Y.G. participated in the challenge under the team name “Hongyang Li and Yuanfang Guan.” C.C. and J.C. participated in the challege under the team names “LiPingChun,” “Guacamole,” and “Lavawizard.” A.H., B.S, G.S., and M.R.C. participated in the challenge under the team names “imp” and “imp1.” A.C., F.G., L.N., M.M., M.J.C., and P.P. participated in the challenge under the team name “BrokenNodes.” C.H. and K.Y.Y. participated in the challenge under the team names “CUImpute1,” “CUWA,” and “ICU.” J.P.S., S.S.B., and Y.S.S. participated in the challenge under the team name “Song Lab.” S.M. and Z.Z. participated in the challenge under the team name “NittanyLions.” W.T., Y.S., Y.S., and Y.S. participated in the challenge under the team name “KKT-ENCODE.”

M.S. and J.A. and R.S. and N.F. and J.H. and K.L. and L.J. and X.Y. and M.C. performed experiments to create the blind test set used to evaluate the methods.

## 4.1.1 Acknowledgements

We would like to thank Alan Min for providing feedback on a draft of the manuscript, Oana Ursu for suggesting analyses, and Jason Ernst for providing manuscript feedback and ChromImpute imputations on the challenge data.

## References

[1] Roadmap Epigenomics Consortium, Anshul Kundaje, Wouter Meuleman, Jason Ernst, Misha Bilenky, Angela Yen, Alireza Heravi-Moussavi, Pouya Kheradpour, Zhizhuo Zhang, Jianrong Wang, Michael J Ziller, Viren Amin, John W Whitaker, Matthew D Schultz, Lucas D Ward, Abhishek Sarkar, Gerald Quon, Richard S Sandstrom, Matthew L Eaton, Yi-Chieh Wu, Andreas R Pfenning, Xinchen Wang, Melina Claussnitzer, Yaping Liu, Cristian Coarfa, R Alan Harris, Noam Shoresh, Charles B Epstein, Elizabeta Gjoneska, Danny Leung, Wei Xie, R David Hawkins, Ryan Lister, Chibo Hong, Philippe Gascard, Andrew J Mungall, Richard Moore, Eric Chuah, Angela Tam, Theresa K Canfield, R Scott Hansen, Rajinder Kaul, Peter J Sabo, Mukul S Bansal, Annaick Carles, Jesse R Dixon, Kai-How Farh, Soheil Feizi, Rosa Karlic, Ah-Ram Kim, Ashwinikumar Kulkarni, Daofeng Li, Rebecca Lowdon, Ginell Elliott, Tim R Mercer, Shane J Neph, Vitor Onuchic, Paz Polak, Nisha Rajagopal, Pradipta Ray, Richard C Sallari, Kyle T Siebenthall, Nicholas A Sinnott-Armstrong, Michael Stevens, Robert E Thurman, Jie Wu, Bo Zhang, Xin Zhou, Arthur E Beaudet, Laurie A Boyer, Philip L De Jager, Peggy J Farnham, Susan J Fisher, David Haussler, Steven J M Jones, Wei Li, Marco A Marra, Michael T McManus, Shamil Sunyaev, James A Thomson, Thea D Tlsty, Li-Huei Tsai, Wei Wang, Robert A Waterland, Michael Q Zhang, Lisa H Chadwick, Bradley E Bernstein, Joseph F Costello, Joseph R Ecker, Martin Hirst, Alexander Meissner, Aleksandar Milosavljevic, Bing Ren, John A Stamatoyannopoulos, Ting Wang, and Manolis Kellis. Integrative analysis of 111 reference human epigenomes. Nature, 518(7539):317–330, February 2015.

[2] ENCODE Project Consortium, Jill E Moore, Michael J Purcaro, Henry E Pratt, Charles B Epstein, Noam Shoresh, Jessika Adrian, Trupti Kawli, Carrie A Davis, Alexander Dobin, Rajinder Kaul, Jessica Halow, Eric L Van Nostrand, Peter Freese, David U Gorkin, Yin Shen, Yupeng He, Mark Mackiewicz, Florencia Pauli-Behn, Brian A Williams, Ali Mortazavi, Cheryl A Keller, Xiao-Ou Zhang, Shaimae I Elhajjajy, Jack Huey, Diane E Dickel, Valentina Snetkova, Xintao Wei, Xiaofeng Wang, Juan Carlos Rivera-Mulia, Joel Rozowsky, Jing Zhang, Surya B Chhetri, Jialing Zhang, Alec Victorsen, Kevin P White, Axel Visel, Gene W Yeo, Christopher B Burge, Eric Lécuyer, David M Gilbert, Job Dekker, John Rinn, Eric M Mendenhall, Joseph R Ecker, Manolis Kellis, Robert J Klein, William S Noble, Anshul Kundaje, Roderic Guigó, Peggy J Farnham, J Michael Cherry, Richard M Myers, Bing Ren, Brenton R Graveley, Mark B Gerstein, Len A Pennacchio, Michael P Snyder, Bradley E Bernstein, Barbara Wold, Ross C Hardison, Thomas R Gingeras, John A Stamatoyannopoulos, and Zhiping Weng. Expanded encyclopaedias of DNA elements in the human and mouse genomes. Nature, 583(7818):699–710, July 2020.

[3] Hendrik G Stunnenberg, International Human Epigenome Consortium, and Martin Hirst. The international human epigenome consortium: A blueprint for scientific collaboration and discovery. Cell, 167(5):1145–1149, November 2016.

[4] Jordan A Ramilowski, Chi Wai Yip, Saumya Agrawal, Jen-Chien Chang, Yari Ciani, Ivan V Kulakovskiy, Mickaël Mendez, Jasmine Li Ching Ooi, John F Ouyang, Nick Parkinson, Andreas Petri, Leonie Roos, Jessica Severin, Kayoko Yasuzawa, Imad Abugessaisa, Altuna Akalin, Ivan V Antonov, Erik Arner, Alessandro Bonetti, Hidemasa Bono, Beatrice Borsari, Frank Brombacher, Christopher J F Cameron, Carlo Vittorio Cannistraci, Ryan Cardenas, Melissa Cardon, Howard Chang, Josée Dostie, Luca Ducoli, Alexander Favorov, Alexandre Fort, Diego Garrido, Noa Gil, Juliette Gimenez, Reto Guler, Lusy Handoko, Jayson Harshbarger, Akira Hasegawa, Yuki Hasegawa, Kosuke Hashimoto, Norihito Hayatsu, Peter Heutink, Tetsuro Hirose, Eddie L Imada, Masayoshi Itoh, Bogumil Kaczkowski, Aditi Kanhere, Emily Kawabata, Hideya Kawaji, Tsugumi Kawashima, S Thomas Kelly, Miki Kojima, Naoto Kondo, Haruhiko Koseki, Tsukasa Kouno, Anton Kratz, Mariola Kurowska-Stolarska, Andrew Tae Jun Kwon, Jeffrey Leek, Andreas Lennartsson, Marina Lizio, Fernando LópezRedondo, Joachim Luginbühl, Shiori Maeda, Vsevolod J Makeev, Luigi Marchionni, Yulia A Medvedeva, Aki Minoda, Ferenc Müller, Manuel Muñoz-Aguirre, Mitsuyoshi Murata, Hiromi Nishiyori, Kazuhiro R Nitta, Shuhei Noguchi, Yukihiko Noro, Ramil Nurtdinov, Yasushi Okazaki, Valerio Orlando, Denis Paquette, Callum J C Parr, Owen J L Rackham, Patrizia Rizzu, Diego Fernando Sánchez Martinez, Albin Sandelin, Pillay Sanjana, Colin A M Semple, Youtaro Shibayama, Divya M Sivaraman, Takahiro Suzuki, Suzannah C Szumowski, Michihira Tagami, Martin S Taylor, Chikashi Terao, Malte Thodberg, Supat Thongjuea, Vidisha Tripathi, Igor Ulitsky, Roberto Verardo, Ilya E Vorontsov, Chinatsu Yamamoto, Robert S Young, J Kenneth Baillie, Alistair R R Forrest, Roderic Guigó, Michael M Hoffman, Chung Chau Hon, Takeya Kasukawa, Sakari Kauppinen, Juha Kere, Boris Lenhard, Claudio Schneider, Harukazu Suzuki, Ken Yagi, Michiel J L de Hoon, Jay W Shin, and Piero Carninci. Functional annotation of human long noncoding RNAs via molecular phenotyping. Genome Res., 30(7):1060–1072, July 2020.

[5] GTEx Consortium, Laboratory, Data Analysis &Coordinating Center (LDACC)—Analysis Working Group, Statistical Methods groups—Analysis Working Group, Enhancing GTEx (eGTEx) groups, NIH Common Fund, NIH/NCI, NIH/NHGRI, NIH/NIMH, NIH/NIDA, Biospecimen Collection Source Site—NDRI, Biospecimen Collection Source Site—RPCI, Biospecimen Core Resource—VARI, Brain Bank Repository—University of Miami Brain Endowment Bank, Leidos Biomedical—Project Management, ELSI Study, Genome Browser Data Integration &Visualization—EBI, Genome Browser Data Integration &Visualization—UCSC Genomics Institute, University of California Santa Cruz, Lead analysts:, Laboratory, Data Analysis &Coordinating Center (LDACC):, NIH program management:, Biospecimen collection:, Pathology:, eQTL manuscript working group:, Alexis Battle, Christopher D Brown, Barbara E Engelhardt, and Stephen B Montgomery. Genetic effects on gene expression across human tissues. Nature, 550(7675):204–213, October 2017.

[6] Rik G H Lindeboom, Aviv Regev, and Sarah A Teichmann. Towards a human cell atlas: Taking notes from the past. Trends Genet., April 2021.

[7] Jason Ernst and Manolis Kellis. Large-scale imputation of epigenomic datasets for systematic annotation of diverse human tissues. Nat. Biotechnol., 33(4):364–376, April 2015.

[8] Wei-Li Guo and De-Shuang Huang. An efficient method to transcription factor binding sites imputation via simultaneous completion of multiple matrices with positional consistency. Mol. Biosyst., 13(9):1827–1837, August 2017.

[9] Qian Qin and Jianxing Feng. Imputation for transcription factor binding predictions based on deep learning. PLoS Comput. Biol., 13(2):e1005403, February 2017.

[10] Timothy J Durham, Maxwell W Libbrecht, J Jeffry Howbert, Jeff Bilmes, and William Stafford Noble. PREDICTD PaRallel epigenomics data imputation with cloud-based tensor decomposition. Nat. Commun., 9(1):1402, April 2018.

[11] Jacob Schreiber, Timothy Durham, Jeffrey Bilmes, and William Stafford Noble. Avocado: a multi-scale deep tensor factorization method learns a latent representation of the human epigenome. Genome Biol., 21(1):81, March 2020.

[12] Jacob Schreiber, Jeffrey Bilmes, and William Stafford Noble. Completing the EN-CODE3 compendium yields accurate imputations across a variety of assays and human biosamples. Genome Biol., 21(1):82, March 2020.

[13] Carles A Boix, Benjamin T James, Yongjin P Park, Wouter Meuleman, and Manolis Kellis. Regulatory genomic circuitry of human disease loci by integrative epigenomics. Nature, 590(7845):300–307, February 2021.

[14] Jennifer Harrow, Adam Frankish, Jose M Gonzalez, Electra Tapanari, Mark Diekhans, Felix Kokocinski, Bronwen L Aken, Daniel Barrell, Amonida Zadissa, Stephen Searle, If Barnes, Alexandra Bignell, Veronika Boychenko, Toby Hunt, Mike Kay, Gaurab Mukherjee, Jeena Rajan, Gloria Despacio-Reyes, Gary Saunders, Charles Steward, Rachel Harte, Michael Lin, Cédric Howald, Andrea Tanzer, Thomas Derrien, Jacqueline Chrast, Nathalie Walters, Suganthi Balasubramanian, Baikang Pei, Michael Tress, Jose Manuel Rodriguez, Iakes Ezkurdia, Jeltje van Baren, Michael Brent, David Haussler, Manolis Kellis, Alfonso Valencia, Alexandre Reymond, Mark Gerstein, Roderic Guigó, and Tim J Hubbard. GENCODE: the reference human genome annotation for the ENCODE project. Genome Res., 22(9):1760–1774, September 2012.

[15] FANTOM Consortium and the Riken PMI and CLST (DGT), Alistair R R Forrest, Hideya Kawaji, Michael Rehli, J Kenneth Baillie, Michiel J L de Hoon, Vanja Haberle, Timo Lassmann, Ivan V Kulakovskiy, Marina Lizio, Masayoshi Itoh, Robin Andersson, Christopher J Mungall, Terrence F Meehan, Sebastian Schmeier, Nicolas Bertin, Mette Jørgensen, Emmanuel Dimont, Erik Arner, Christian Schmidl, Ulf Schaefer, Yulia A Medvedeva, Charles Plessy, Morana Vitezic, Jessica Severin, Colin A Semple, Yuri Ishizu, Robert S Young, Margherita Francescatto, Intikhab Alam, Davide Albanese, Gabriel M Altschuler, Takahiro Arakawa, John A C Archer, Peter Arner, Magda Babina, Sarah Rennie, Piotr J Balwierz, Anthony G Beckhouse, Swati Pradhan-Bhatt, Judith A Blake, Antje Blumenthal, Beatrice Bodega, Alessandro Bonetti, James Briggs, Frank Brombacher, A Maxwell Burroughs, Andrea Califano, Carlo V Cannistraci, Daniel Carbajo, Yun Chen, Marco Chierici, Yari Ciani, Hans C Clevers, Emiliano Dalla, Carrie A Davis, Michael Detmar, Alexander D Diehl, Taeko Dohi, Finn Drabløs, Albert S B Edge, Matthias Edinger, Karl Ekwall, Mitsuhiro Endoh, Hideki Enomoto, Michela Fagiolini, Lynsey Fairbairn, Hai Fang, Mary C Farach-Carson, Geoffrey J Faulkner, Alexander V Favorov, Malcolm E Fisher, Martin C Frith, Rie Fujita, Shiro Fukuda, Cesare Furlanello, Masaaki Furino, Jun-Ichi Furusawa, Teunis B Geijtenbeek, Andrew P Gibson, Thomas Gingeras, Daniel Goldowitz, Julian Gough, Sven Guhl, Reto Guler, Stefano Gustincich, Thomas J Ha, Masahide Hamaguchi, Mitsuko Hara, Matthias Harbers, Jayson Harshbarger, Akira Hasegawa, Yuki Hasegawa, Takehiro Hashimoto, Meenhard Herlyn, Kelly J Hitchens, Shannan J Ho Sui, Oliver M Hofmann, Ilka Hoof, Furni Hori, Lukasz Huminiecki, Kei Iida, Tomokatsu Ikawa, Boris R Jankovic, Hui Jia, Anagha Joshi, Giuseppe Jurman, Bogumil Kaczkowski, Chieko Kai, Kaoru Kaida, Ai Kaiho, Kazuhiro Kajiyama, Mutsumi Kanamori-Katayama, Artem S Kasianov, Takeya Kasukawa, Shintaro Katayama, Sachi Kato, Shuji Kawaguchi, Hiroshi Kawamoto, Yuki I Kawamura, Tsugumi Kawashima, Judith S Kempfle, Tony J Kenna, Juha Kere, Levon M Khachigian, Toshio Kitamura, S Peter Klinken, Alan J Knox, Miki Kojima, Soichi Kojima, Naoto Kondo, Haruhiko Koseki, Shigeo Koyasu, Sarah Krampitz, Atsutaka Kubosaki, Andrew T Kwon, Jeroen F J Laros, Weonju Lee, Andreas Lennartsson, Kang Li, Berit Lilje, Leonard Lipovich, Alan Mackay-Sim, RiIchiroh Manabe, Jessica C Mar, Benoit Marchand, Anthony Mathelier, Niklas Mejhert, Alison Meynert, Yosuke Mizuno, David A de Lima Morais, Hiromasa Morikawa, Mitsuru Morimoto, Kazuyo Moro, Efthymios Motakis, Hozumi Motohashi, Christine L Mummery, Mitsuyoshi Murata, Sayaka Nagao-Sato, Yutaka Nakachi, Fumio Nakahara, Toshiyuki Nakamura, Yukio Nakamura, Kenichi Nakazato, Erik van Nimwegen, Noriko Ninomiya, Hiromi Nishiyori, Shohei Noma, Shohei Noma, Tadasuke Noazaki, Soichi Ogishima, Naganari Ohkura, Hiroko Ohimiya, Hiroshi Ohno, Mitsuhiro Ohshima, Mariko Okada-Hatakeyama, Yasushi Okazaki, Valerio Orlando, Dmitry A Ovchinnikov, Arnab Pain, Robert Passier, Margaret Patrikakis, Helena Persson, Silvano Piazza, James G D Prendergast, Owen J L Rackham, Jordan A Ramilowski, Mamoon Rashid, Timothy Ravasi, Patrizia Rizzu, Marco Roncador, Sugata Roy, Morten B Rye, Eri Saijyo, Antti Sajantila, Akiko Saka, Shimon Sakaguchi, Mizuho Sakai, Hiroki Sato, Suzana Savvi, Alka Saxena, Claudio Schneider, Erik A Schultes, Gundula G Schulze-Tanzil, Anita Schwegmann, Thierry Sengstag, Guojun Sheng, Hisashi Shimoji, Yishai Shimoni, Jay W Shin, Christophe Simon, Daisuke Sugiyama, Takaai Sugiyama, Masanori Suzuki, Naoko Suzuki, Rolf K Swoboda, Peter A C ‘t Hoen, Michihira Tagami, Naoko Takahashi, Jun Takai, Hiroshi Tanaka, Hideki Tatsukawa, Zuotian Tatum, Mark Thompson, Hiroo Toyodo, Tetsuro Toyoda, Elvind Valen, Marc van de Wetering, Linda M van den Berg, Roberto Verado, Dipti Vijayan, Ilya E Vorontsov, Wyeth W Wasserman, Shoko Watanabe, Christine A Wells, Louise N Winteringham, Ernst Wolvetang, Emily J Wood, Yoko Yamaguchi, Masayuki Yamamoto, Misako Yoneda, Yohei Yonekura, Shigehiro Yoshida, Susan E Zabierowski, Peter G Zhang, Xiaobei Zhao, Silvia Zucchelli, Kim M Summers, Harukazu Suzuki, Carsten O Daub, Jun Kawai, Peter Heutink, Winston Hide, Tom C Freeman, Boris Lenhard, Vladimir B Bajic, Martin S Taylor, Vsevolod J Makeev, Albin Sandelin, David A Hume, Piero Carninci, and Yoshihide Hayashizaki. A promoter-level mammalian expression atlas. Nature, 507(7493):462– 470, March 2014.

[16] Sage Bionetworks. [no title]. https://www.synapse.org/!Synapse:syn6131484/wiki/. Accessed: 2021-5-12.

[17] Jacob Schreiber, Ritambhara Singh, Jeffrey Bilmes, and William Stafford Noble. A pitfall for machine learning methods aiming to predict across cell types. Genome Biol., 21(1):282, November 2020.

[18] Yong Zhang, Tao Liu, Clifford A Meyer, Jérôme Eeckhoute, David S Johnson, Bradley E Bernstein, Chad Nusbaum, Richard M Myers, Myles Brown, Wei Li, and X Shirley Liu. Model-based analysis of ChIP-Seq (MACS). Genome Biol., 9(9):R137, September 2008.

[19] Jinwook Lee, Daniel Kim, Grey Cristoforo, Chuan-Sheng Foo, Chris Probert, Nathan Beley, and Anshul Kundaje. ENCODE ATAC-seq pipeline, December 2019.

[20] Ben Langmead and Steven L Salzberg. Fast gapped-read alignment with bowtie 2. Nat. Methods, 9(4):357–359, March 2012.

[21] Aaron McKenna, Matthew Hanna, Eric Banks, Andrey Sivachenko, Kristian Cibulskis, Andrew Kernytsky, Kiran Garimella, David Altshuler, Stacey Gabriel, Mark Daly, and Mark A DePristo. The genome analysis toolkit: a MapReduce framework for analyzing next-generation DNA sequencing data. Genome Res., 20(9):1297–1303, September 2010.

[22] Haley M Amemiya, Anshul Kundaje, and Alan P Boyle. The ENCODE blacklist: Identification of problematic regions of the genome. Sci. Rep., 9(1):9354, June 2019.

[23] Jin Lee, J Seth Strattan annashcherbina, Karl Sebby, Meenakshi Kagda, and Paul L Maurizio. ENCODE-DCC/chip-seq-pipeline2: v1.9.0, May 2021.

[24] Heng Li and Richard Durbin. Fast and accurate short read alignment with Burrows-Wheeler transform. Bioinformatics, 25(14):1754–1760, July 2009.

[25] Yaxing Zhao, Limsoon Wong, and Wilson Wen Bin Goh. How to do quantile normalization correctly for gene expression data analyses. Sci. Rep., 10(1):15534, September 2020.

[26] F William Townes and Rafael A Irizarry. Quantile normalization of single-cell RNA-seq read counts without unique molecular identifiers. Genome Biol., 21(1):160, July 2020.

[27] Nicolas Bonhoure, Gergana Bounova, David Bernasconi, Viviane Praz, Fabienne Lammers, Donatella Canella, Ian M Willis, Winship Herr, Nouria Hernandez, Mauro Delorenzi, and CycliX Consortium. Quantifying ChIP-seq data: a spiking method providing an internal reference for sample-to-sample normalization. Genome Res., 24(7):1157– 1168, July 2014.

[28] Žiga Avsec, Melanie Weilert, Avanti Shrikumar, Sabrina Krueger, Amr Alexandari, Khyati Dalal, Robin Fropf, Charles McAnany, Julien Gagneur, Anshul Kundaje, and Julia Zeitlinger. Base-resolution models of transcription-factor binding reveal soft motif syntax. Nat. Genet., 53(3):354–366, March 2021.

