## Additional File 2 for "The ENCODE Imputation Challenge: A critical assessment of methods for cross-cell type imputation of epigenomic profiles"

### Additional File 2: Supplementary Figures

July 30, 2022

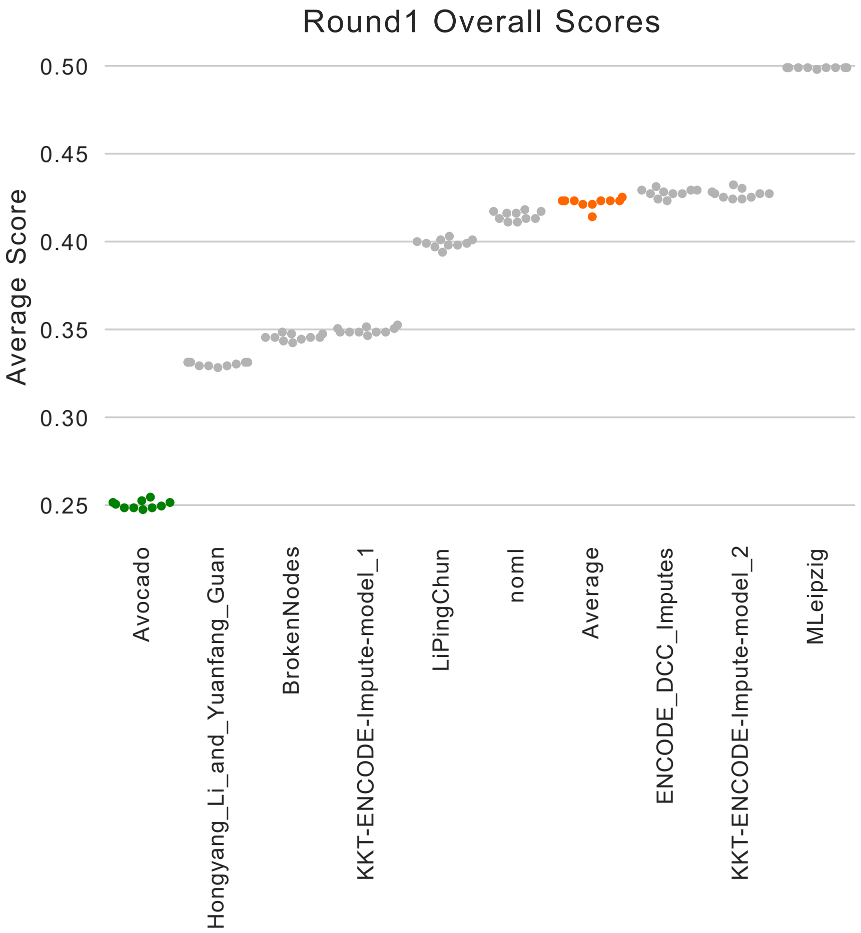


Figure S1: **Round 1 Scores.** The final leaderboard results for each participant in round 1 of the challenge. Each dot is the score calculated on one of ten bootstraps, and the participants are ordered by overall score. Lower scores indicate better performance.

| | team | track.rank | final.rank | teamname |
|---|---|---|---|---|
| *1* | 3393580 | 6.568627 | 1 | Song Lab 3 |
| *2* | 3388185 | 6.725490 | 2 | LiPingChun |
| *3* | 3393847 | 7.294118 | 3 | Guacamole |
| *4* | 3393817 | 7.784314 | 4 | BrokenNodes_v3 |
| *5* | 3393574 | 7.843137 | 5 | Lavawizard |
| *6* | 3393860 | 7.882353 | 6 | CUImpute1 |
| *7* | 3393861 | 8.803922 | 7 | ICU |
| *8* | 3393458 | 10.294118 | 8 | CUWA |
| *9* | 3389318 | 10.490196 | 9 | Song Lab |
| *10* | 3330254 | 10.627451 | 10 | HLYG |
| *11* | 3393417 | 10.921569 | 11 | HLYG v1 |
| *12* | 3393457 | 11.294118 | 12 | imp |
| *13* | 200 | 11.568627 | 13 | ChromImpute |
| *14* | 3393756 | 11.627451 | 14 | imp1 |
| *15* | 3393606 | 11.882353 | 15 | BrokenNodes_v2 |
| *16* | 3379312 | 13.529412 | 16 | CostaLab |
| *17* | 3379072 | 13.549020 | 17 | BrokenNodes |
| *18* | 3393418 | 14.705882 | 18 | HLYG v2 |
| *19* | 100 | 14.862745 | 19 | Average |
| *20* | 0 | 17.078431 | 20 | Avocado |
| *21* | 3393128 | 19.666667 | 21 | Aug2019Imputation |
| *22* | 3393851 | 21.156863 | 22 | NittanyLions2 |
| *23* | 3344979 | 21.666667 | 23 | NittanyLions |
| *24* | 3393579 | 23.215686 | 24 | Song Lab 2 |
| *25* | 3391272 | 24.320000 | 25 | UIOWA Michaelson |
| *26* | 3386902 | 25.607843 | 26 | KKT−ENCODE−Impute |

Figure S2: **Overall ranking using six measures from ChromImpute.** A re-ranking of the methods used in the challenge, including ChromImpute as a baseline, using the six of the relative performance measures used to evaluate the original ChromImpute model. Ranks were computed by ranking team performance for each blind track and metric combination first, then averaging ranks and re-ranking at the track level, and finally averaging to rank overall team performance. Performance measures included (i) genome-wide correlation (GWcorr), (ii) overlap of the top 1% of imputed and observed 25 bp bins, (iii) percentage of top 1% of imputed bins in top 5% of observed bins and (iv) vice versa, and the area under the receiver operating characteristic curve (AUROC) for predicting the (v) top 1% of observed bins with imputed signal or (vi) top 1% of imputed bins using the observed signal.

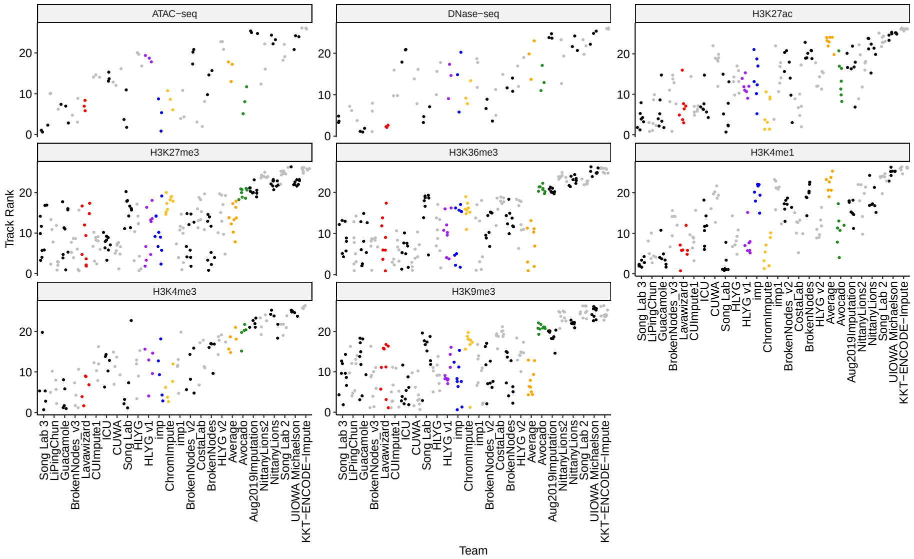


Figure S3: **Rankings per assay using six measures from ChromImpute.** A re-ranking of the methods used in the challenge broken down by assay, with each dot representing an experiment in the blind test set. Colors alternate black and grey by team except for colors that highlight top performing teams as in the main text. ChromImpute and the average predictor are colored in yellow and Avocado in green.

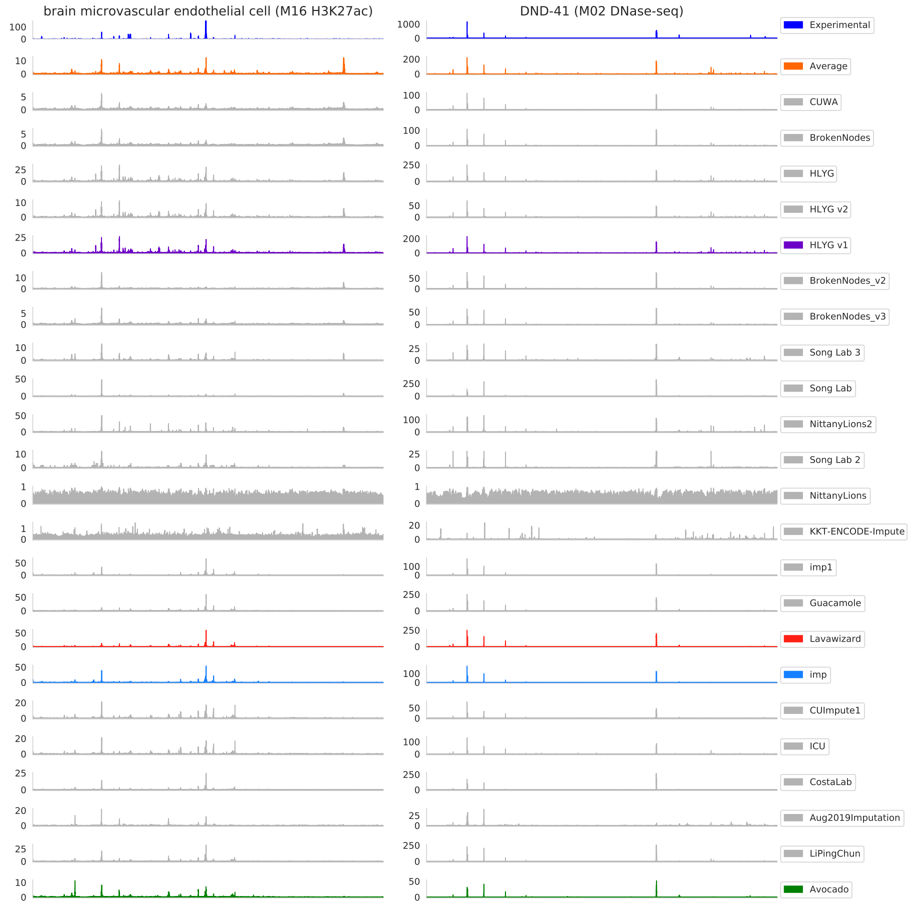


Figure S4: **Imputations from all models at two illustrative loci.** Imputations and experimental signal for H3K27ac in brain microvascular endothelial cells and for DNase-seq in DND-41 cells. The experimental signal is proceeded by all methods, ranked by the maximum of their correlation with the average activity and with the Avocado imputations. The two baseline methods, and the winning submissions, are colored.

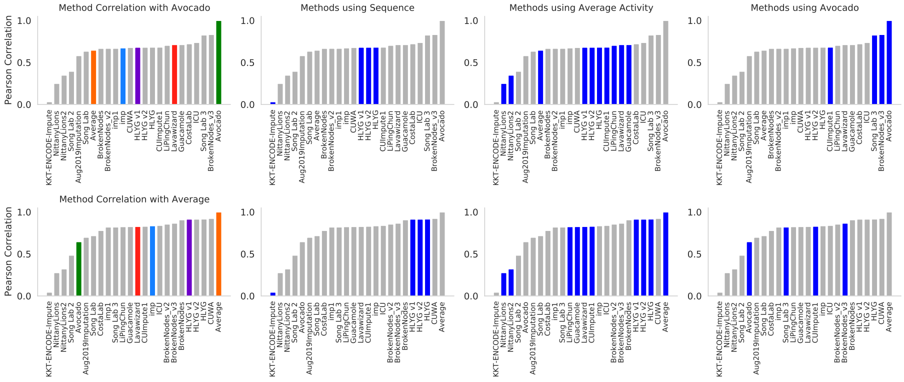


Figure S5: **Method's correlations with the average activity and Avocado** The Pearson correlation of each method with Avocado's imputations (top row) and with the average activity (bottom row). The left column shows the baseline methods and challenge winners colored, ordered by their correlation with either Avocado or the average activity, respectively. The other panels show the same results as the left panel but color methods that use sequence features, the average activity, or Avocado's imputations, as input.

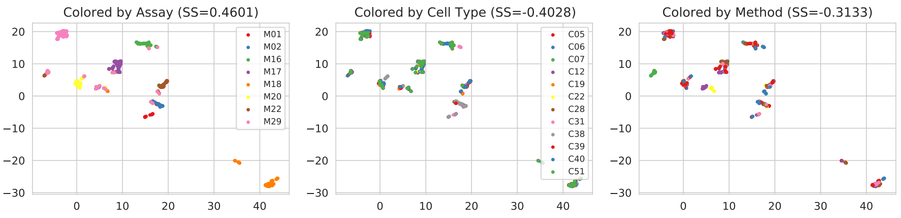


Figure S6: **A projection of all experiments.** A UMAP projection of the experimental and imputed tracks from all teams based on Pearson correlation averaged across each chromosome. The panels are colored, and show the Silhoutte Score when clustering, by assay type, cell type, and method.

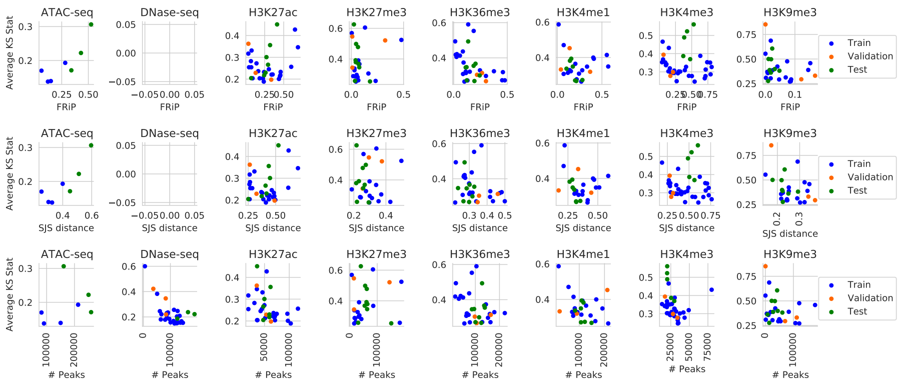


Figure S7: **Quality control metrics for challenge experiments.** Each row displays a different quality control metric, each column displays that measure within a certain assay, and each dot in the panel corresponds to the metric score for a particular experiment. Each dot is colored based on if it is a training (blue), validation (orange), or test set (green) experiment.

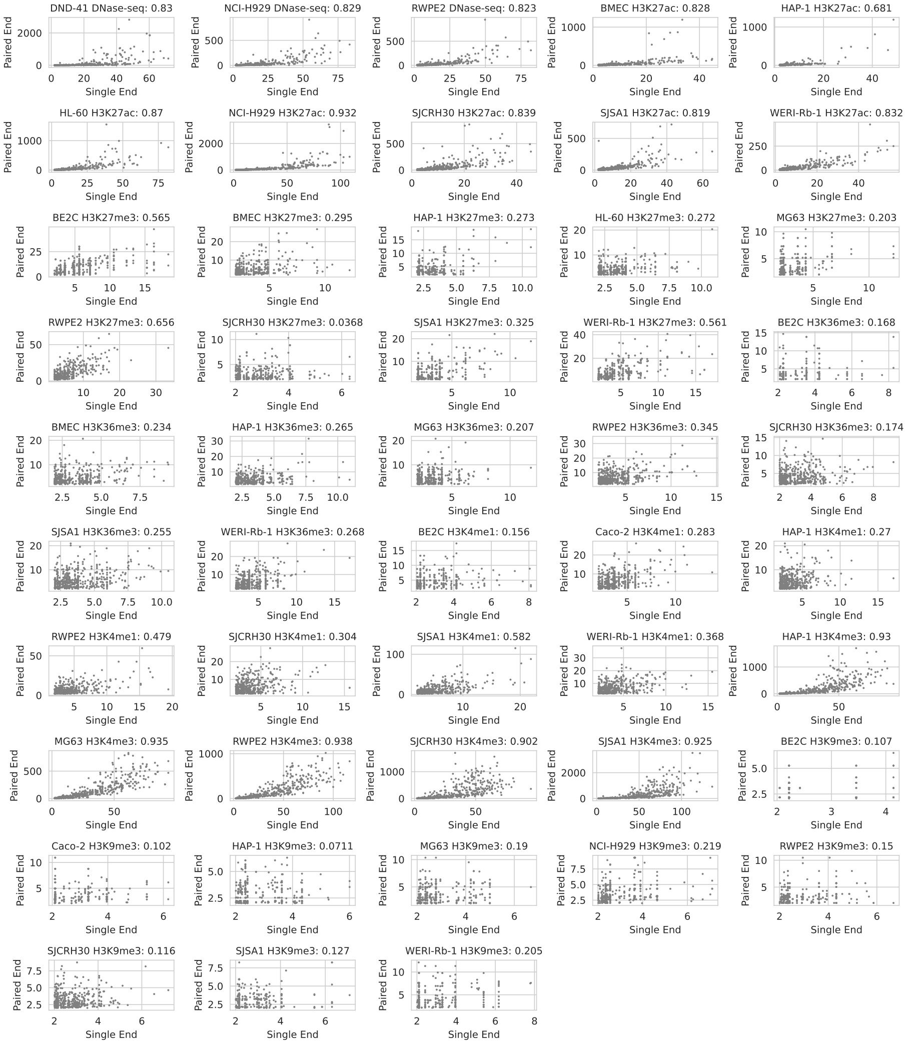


Figure S8: **Signal values with paired-end and single-end processing** For each of the 48 reprocessed experiments, 10Mb in the center of chr1 are considered when the reads are processed using the original paired-end processing or the single-end reprocessing. Only 500 random positions within this block are visualized for each experiment, but the Spearman correlation shown for each panel is calculated across the entire 10Mb block.

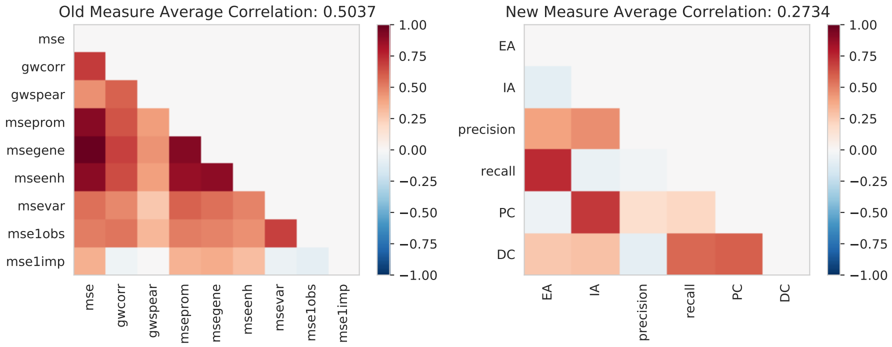


Figure S9: **Rank correlation of performance measures** Rank correlations are calculated between each performance measure by calculating the Spearman correlation across all teams for each pair of measures. For the original nine measures (left), the rank of each team for each measure is averaged across all ten folds. For the new six measures the Spearman correlation is calculated directly on the performance values because there are no bootstraps.

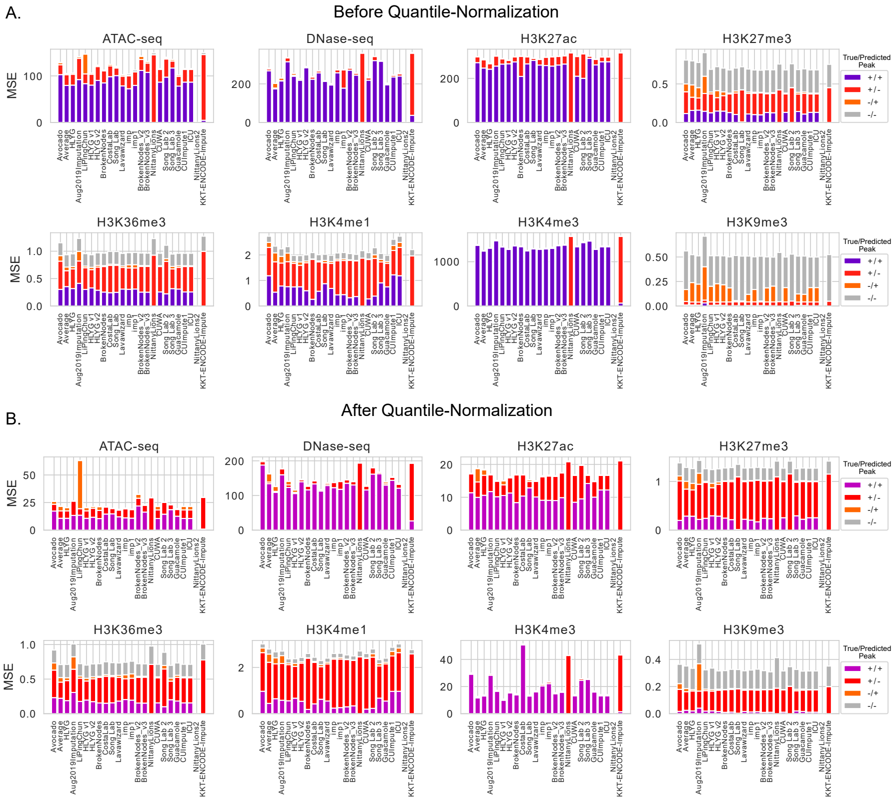


Figure S10: **Error partitioned by peak calls** The genome-wide squared error was partitioned into four categories for each assay: within correctly predicted peaks (+/+), within true peaks incorrectly not predicted to be peaks (+/-), within background regions incorrectly predicted to be peaks (-/+), and within background regions correctly predicted to be background (-/-). True peaks were identified using the MACS2 peak caller and predicted peaks were identified as a predicted signal value $\geq 2$. (A) The error for each method on the original signal. (B) The error for each method on the quantile-normalized signal.
